# High throughput competitive fluorescence polarization assay reveals functional redundancy in the S100 protein family

**DOI:** 10.1101/718155

**Authors:** Márton A. Simon, Péter Ecsédi, Gábor M. Kovács, Ádám L. Póti, Attila Reményi, József Kardos, Gergő Gógl, László Nyitray

## Abstract

The calcium-binding, vertebrate-specific S100 protein family consists of 20 paralogs in humans (referred as the S100ome), with several clinically important members. To assess their interactome, high-throughput, systematic analysis is indispensable, which allows one to get not only qualitative but quantitative insight into their protein-protein interactions (PPIs). We have chosen an unbiased assay, fluorescence polarization (FP) that revealed a partial functional redundancy when the complete S100ome (n=20) was tested against numerous model partners (n=13). Based on their specificity, the S100ome can be grouped into two distinct classes: promiscuous and orphan. In the first group, members bound to several ligands (>4-5) with comparable high affinity, while in the second one, the paralogs bound only one partner weakly, or no ligand was identified (orphan). Our results demonstrate that *in vitro* FP assays are highly suitable for quantitative ligand binding studies of selected protein families. Moreover, we provide evidence that PPI-based phenotypic characterization can complement the information obtained from the sequence-based phylogenetic analysis of the S100ome, an evolutionary young protein family.

**Author summary:** Functional similarity among a protein family can be essential in order to understand proteomic data, to find biomarkers, or in inhibitor design. Proteins with similar functions can compensate the loss-of-function of the others, their expression can co-vary under pathological conditions, and simultaneous targeting can lead to better results in the clinics. To investigate this property one can use sequence-based approaches. However, this path can be difficult. In the case of the vertebrate specific, evolutionary young, S100 family, phylogenetic approaches lead to ambiguous results. To overcome this problem, we applied a high-throughput biochemical approach to experimentally measure the binding affinities of a large number of S100 interactions. We performed unbiased fluorescence polarization assay, involving the complete human S100ome (20 paralogs) and 13 known interaction partners. We used this measured 20×13 (260) protein-protein interaction array to reveal the functional relationships within the family. Our work provide a general framework for studies focusing on phenotype-based domain classification.

## Introduction

Biochemical characterization of protein-protein interactions (PPIs) is a challenging field in molecular life sciences, which is usually limited to the determination of steady state dissociation constants [1]. The accurate determination of thermodynamic parameters of molecular interactions is performed by fast, but superficial high-throughput (HTP) methods. In the literature several HTP approaches are applied such as co-immunoprecipitation [2], yeast two hybrid and spot assays [3], pull-down assay [4], holdup assay [5] and direct fluorescence polarization/anisotropy [6]. In direct fluorescence polarization (FP) experiments, a fluorescent probe (usually a labeled peptide) is titrated with a globular partner. Their association is monitored by the polarization of the emitted light of the fluorophore (Fig 1A). In a modified FP experiment called competitive assay, both the probe and partner concentration are fixed, and the reaction mixture is titrated with an unlabeled competitor molecule (peptide or protein). Depolarization of the emitted light is indicative of the competition between the probe and the competitor in binding to the partner (Fig 1BC). While direct FP can be perturbed by the presence of the fluorescent dye, the competitive assay is unbiased and therefore more suitable for accurate HTP measurements of dissociation constants [7,8].

**Fig 1.**
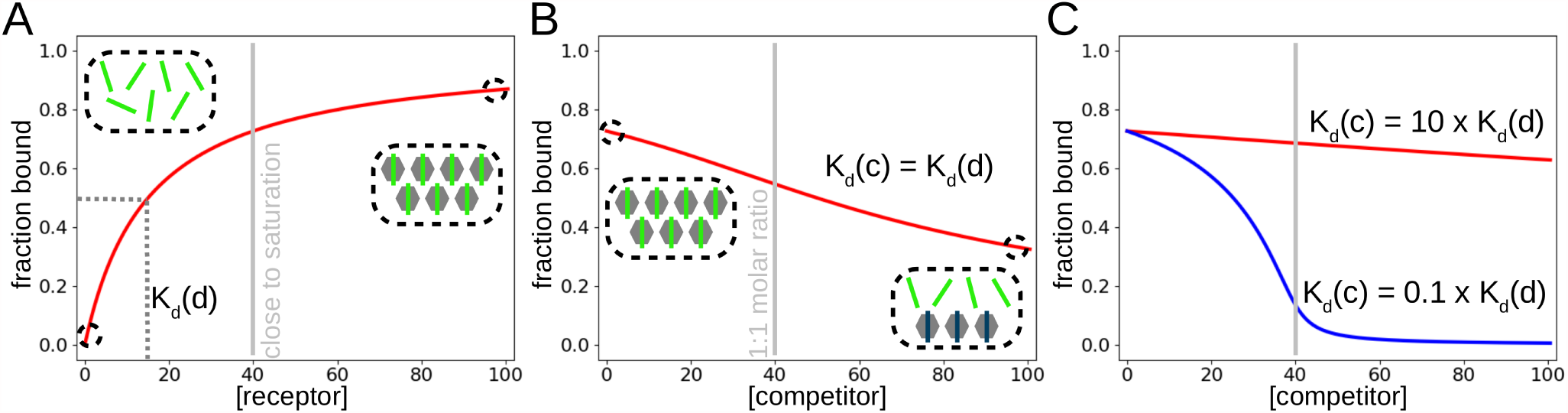
**A:** Fluorescence polarization/anisotropy experiments can be performed with direct and competitive titrations. In direct assay, the concentration of the protein of interest is increased in the presence of tracer amount of labeled peptide. Upon complex formation, the hydrodynamic radius of the tracer increases causing slower rotation and therefore lower depolarization of the emitted light. In the direct assay, one can measure the minimal and maximal polarization values, a dissociation constant and importantly, an optimal concentration can be easily determined for competitive assays, which is usually the concentration corresponding to 60-80% saturation. **B:** in a competitive assay, the concentration of the protein of interest is set to this concentration and one can titrate the reaction mixture with a competitor. The competition results in increased level of free labeled peptide and consequently high depolarization of the emitted light. **C:** competitive FP is not affected by the presence of a labeling group in the peptide (unbiased) and has a high dynamic range (approximately 2 orders of magnitudes around the dissociation constant of the probe). At high concentrations, it can be also used to determine the stoichiometry of the interaction for strong interactions.

S100 proteins belong to the superfamily of EF-hand containing calcium-binding proteins. They appeared in early vertebrates and consist of 20 core paralogs in the human proteome [9]. S100s are associated with several disease conditions, such as cardiomyopathies, cancer, inflammatory and neurodegenerative diseases, in which overexpression of S100 proteins can be observed in the affected cells [10–12]. Due to this reason, they are emerging bio-markers and also promising therapeutic targets [13]. Despite their growing importance, the literature still lacks their comprehensive and systematic analysis, which would be essential for developing rational strategies for drug development. Similarly to calmodulin, they can interact with protein or peptide targets in a calcium-dependent manner [14]. They are generally considered as relatively low specificity proteins, with dozens of interaction partners, among them they are unable to maintain a high selectivity [15]. In this study we determined the interaction profile of the full human S100 family (termed here as the S100ome) against a set of diverse known S100 partners (and some of their paralogs) systematically, including kinases such as RSK1 [16] and its paralogs MK2 and MNK1; cytoskeletal elements such as CapZ [17] (commonly known as TRTK12), NMIIA [18], ezrin [19], FOR20 and its paralog FOP [20]; membrane proteins such as NCX1 [15] and TRPM4 [21]; and other signaling proteins such as the tumor suppressor p53 [22–24], SIP [15] and MDM4 [23].

## Results

### Mapping the S100ome with FP measurements

The interactions between S100 homodimers and their selected labeled peptide partners were studied first by direct FP assay (Fig S1-13.). We have found that all reasonable S100 interactions gave an experimental window of 50-200 mP (polarization). If significant binding was detected (Kd < 200 µM) between a labeled peptide and an S100 protein, a subsequent competitive FP assay was performed. In cases, where no labeled peptide was available (e.g. when globular protein domains were used as competitors), we used non-cognate tracers against all possible S100 proteins. Additionally, we tested the possible binding between these competitors and the non-cognate probes in direct FP experiments to eliminate the possibility of re-binding (Fig S14.). This way, we tested 180 unique direct and 150 unique competitive interactions and found 89, and 66 significant interactions, respectively (Table 2., Fig S1-13.).

**Table 1.**
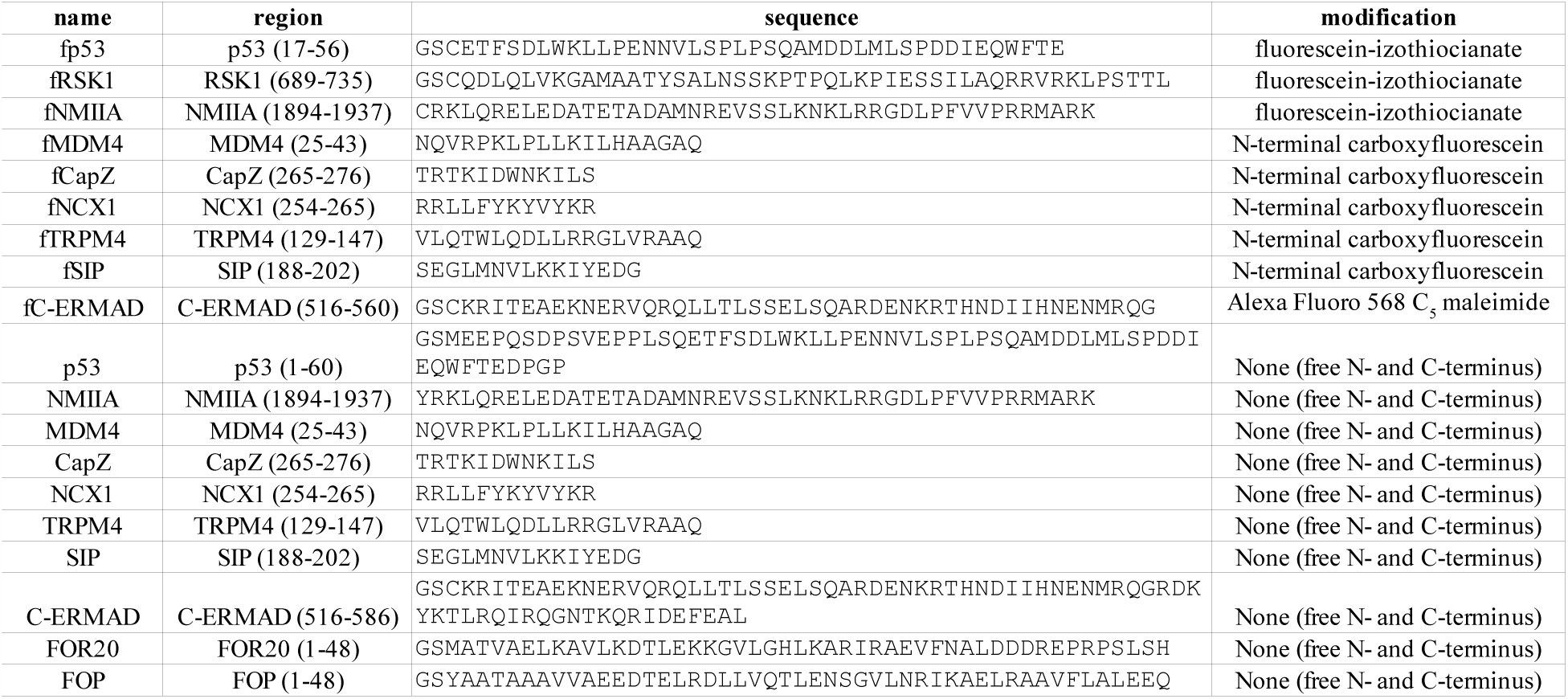
Peptides used in this study

**Table 2.**
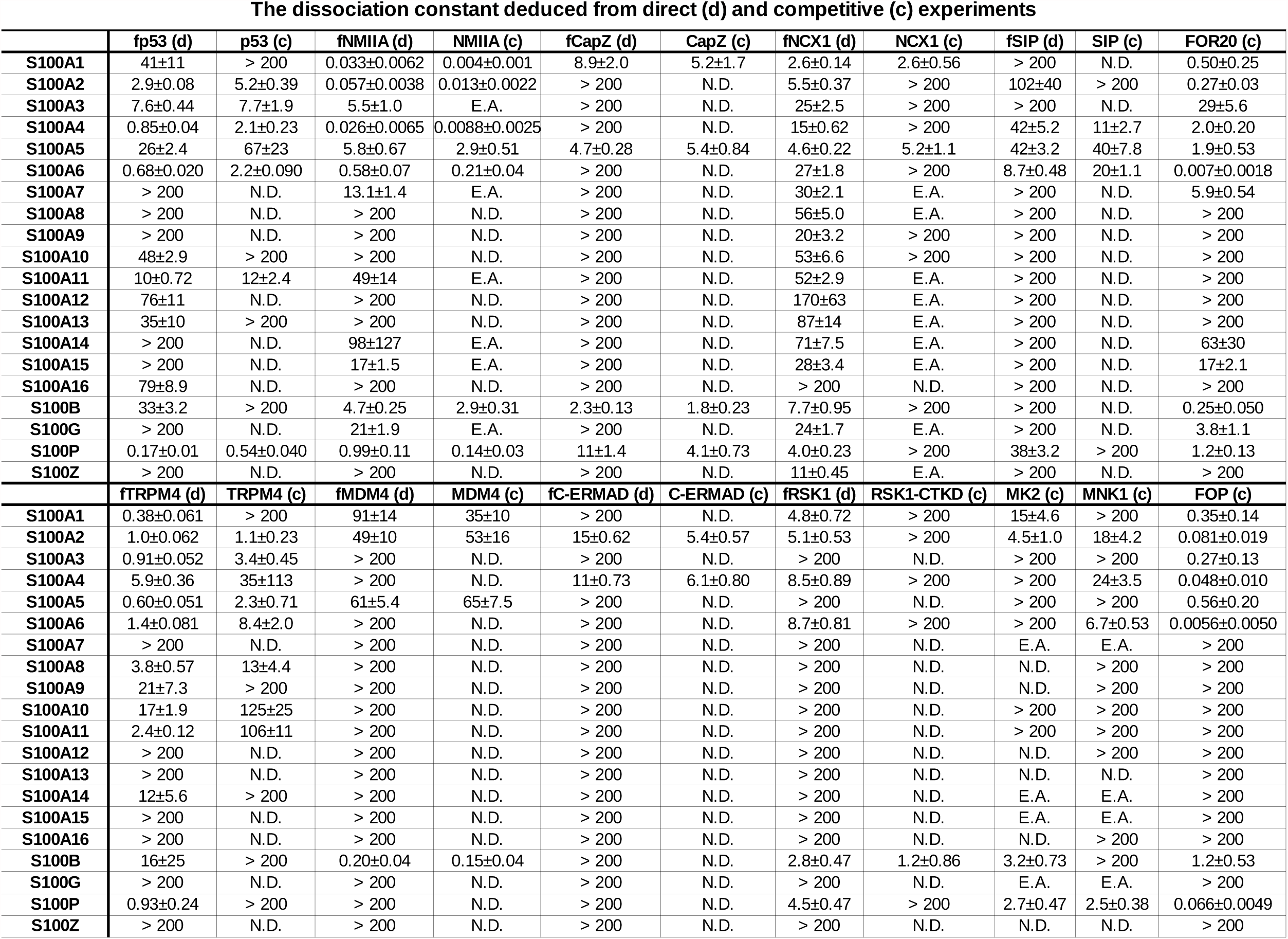
Quantitative characterization of interactions between S100 proteins and their selected partners by FP (N.D.: not determined; E.A.: experimental artifact).

As mentioned previously, competitive FP provides unbiased (or more specific) affinities, unaffected by the chemical labeling, making it a better tool to measure protein-protein interactions (Fig 2AB). Nevertheless, there are some pitfalls (Fig 2.), which should be taken into consideration while analyzing competitive data. First of all, the experimental window of the competitive measurement should be the same as the experimental window of the direct measurement (Fig 2C). Studying large biomolecules (e.g. globular proteins) in a competitive experiment often results in an increased base polarization (P_min_) due to the change in biophysical properties of the reaction mixture (e.g. change in viscosity). Moreover, during a competition experiment, it is possible that the competitor can interact with the probe itself, which can also cause an increase in the base polarization (Fig S14.). In rare cases, saturation polarization can be also altered. Additionally, experimental artifacts of unknown origin can be observed occasionally (Fig 2D). Here, a sharp decline can be detected during the titration, which results in an IC50 value smaller than the fixed receptor concentration. This observed sub-stoichiometric complex formation should be handled with extra care as it is likely due to unexpected biophysical phenomena, such as protein aggregation. To standardize and automatize data handling and to eliminate subjective factors, we developed a Python-based universal program, called ProFit for fitting all direct and competitive experimental data (freely available at https://github.com/GoglG/ProFit).

**Fig 2.**
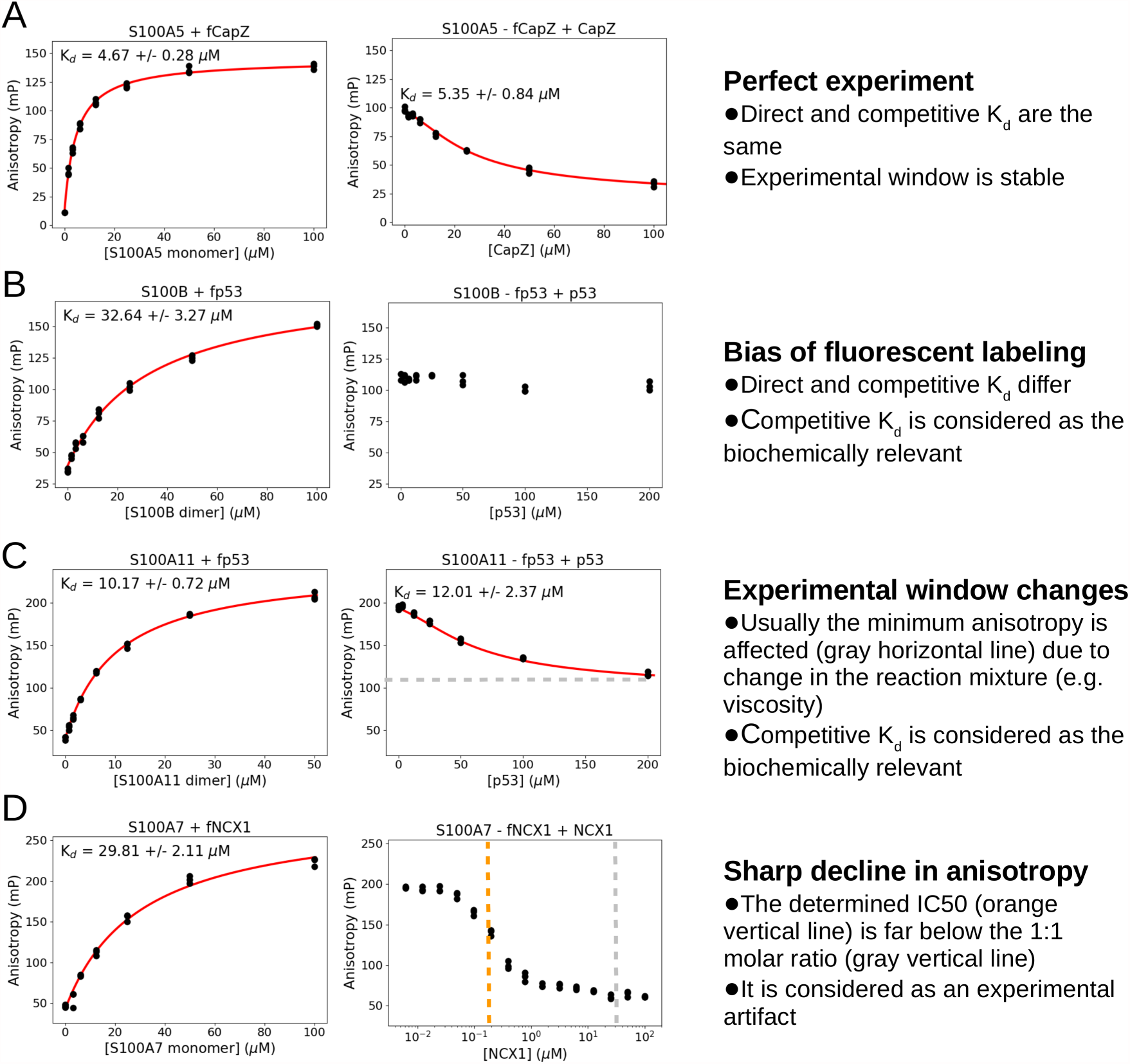
**A:** in “perfect experiments”, the experimental window is stable and the dissociation constants match between the (cognate) probe and the competitor. **B:** as often occurs, fluorescent labeling can alter the binding affinity, resulting in false positive interaction partners in direct FP experiments. In other cases, the effect is softer and it only causes a dimming effect on the biochemical constant. **C:** the reliable experimental window can be different in a competitive experiment. If the change is not extreme, the competitive K_d_ can be considered (with caution) as the relevant biochemical constant. **D:** in some cases, a rapid decline can be observed in the polarization. In this case, the experimentally determined IC50 value should not be used as a dissociation constant. This phenomena can be explained by a competitor induced biophysical transition, e.g. aggregation or precipitation. In this final case, it is very important to redetermine the concentrations of the receptor and the competitor and to repeat the experiment at different receptor concentrations to properly discriminate the stoichiometric molar ratio from the observed IC50 value.

### Validation with ITC measurements

The biochemically described S100 binding motifs, found in the literature, show an extremely low sequence similarity [15,23] (Fig 3A). Mostly linear segments are recognized by the human S100ome, however, no consensus S100 binding sequence can be defined[15]. In general, hydrophobic residues are preferred, but additional basic residues can also be favored in some instances. Moreover, S100 proteins can form two types of complexes (Fig 3B). Earlier studies showed that a symmetric S100 dimer can recognize two identical binding motifs, symmetrically [17,25,26]. In recent studies however, several asymmetric complexes were also described [18,27,28]. In those cases, an S100 dimer captures a single straddling the two binding sites. As the binding affinity highly depends on the stoichiometry of the interaction, we selected a set of significant, peptide-based interactions for isothermal titration calorimetric (ITC) measurements. This way, we validated the interactions that were originally detected by the FP assay and determined the binding stoichiometry in all instances.

**Fig 3.**
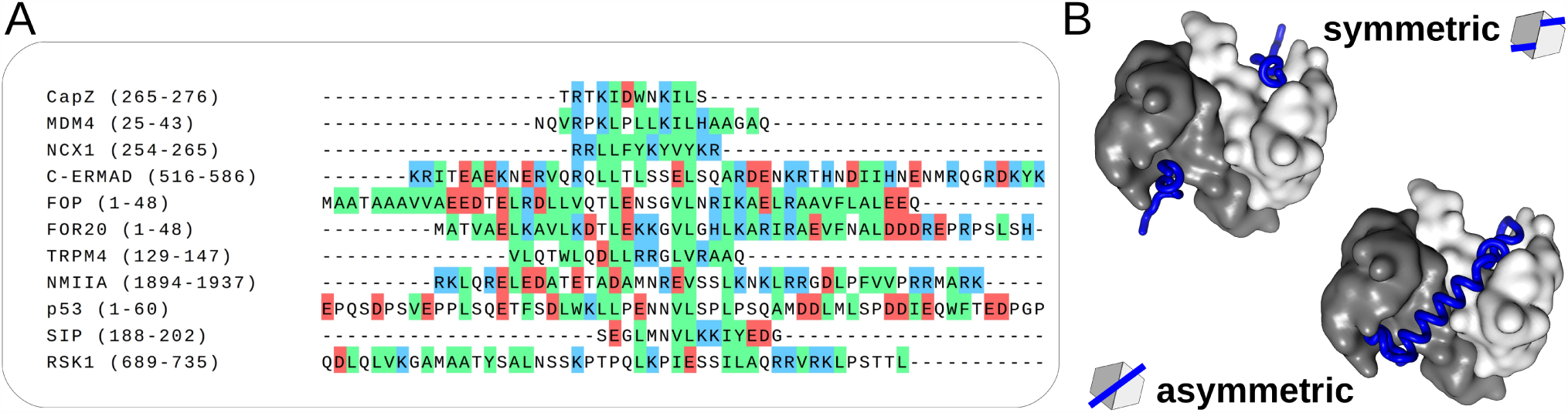
**A:** multiple short linear motifs are recognized by S100 family members, however no consensus binding motif can be defined for the protein family, as indicated here by the sequence alignment of several S100 binding motifs. Though it is noteworthy that hydrophobic residues (green) are preferred, basic residues are also welcome in some cases. **B:** S100 proteins act as dimers and are capable of interacting in two distinct ways with other proteins. On the left, a symmetric complex is shown, where one S100 dimer interacts with two peptides (S100B-CapZ, [17]). By contrast, a single interacting partner can bind to one S100 dimer asymmetrically, as it is shown on the right side (S100A4-NMIIA, [18]).

All determined K_d_ values correlated well with the data provided by the orthogonal FP measurements (Table 3., Fig S15.). Symmetric interactions were found with CapZ, NCX1, SIP, TRPM4 and MDM4. In cases of CapZ and MDM4, the experimental data was fitted by a two binding site model indicating slightly different affinities and a complex relationship between the S100 monomers. In contrast, asymmetrical interactions were detected with p53, RSK1, C-ERMAD, NMIIA and FOP. These findings confirmed the expected binding stoichiometry in all cases and clarified the binding mode of TRPM4 and FOP. We hypothesize that the binding mode of close paralogs should be identical (symmetric or asymmetric), therefore, asymmetric binding was assumed for MNK1, MK2 (based on RSK1) and FOR20 (based on FOP). We performed these ITC measurements in parallel with the FP experiments and based on the refined stoichiometry, monomer or dimer S100 concentrations were used during the FP data evaluation.

**Table 3.** Quantitative characterization of interactions between S100 proteins and their selected partners by ITC.

| measurement | T / K | N | Reference N | $K_d$ / μM | $\Delta H$ / (kJ/mol) | $-T\Delta S$ (kJ/mol) |
| --- | --- | --- | --- | --- | --- | --- |
| S100A6-FOP | 310 | $0.44 \pm 0.002$ | Previously unknown | $0.088 \pm 0.0073$ | $-73 \pm 0.58$ | 31 |
| S100B-CapZ | 310 | $0.44 \pm 0.002$ | 1 [17] | $3.9 \pm 0.39$ | $-15 \pm 0.37$ | -17 |
| | | $0.343 \pm 0.002$ | | $0.94 \pm 0.03$ | $-3.3 \pm 0.67$ | -33 |
| S100A1-NCX1 | 310 | $1.1 \pm 0.013$ | 1 [15]* | $6.4 \pm 0.58$ | $-35 \pm 0.80$ | 4.0 |
| S100A5-TRPM4 | 310 | $0.89 \pm 0.0037$ | Previously unknown | $1.4 \pm 0.089$ | $-24 \pm 0.21$ | -10 |
| C-ERMAD-S100A4 | 310 | $1.9 \pm 0.055$ | 1.86 [19] | $35 \pm 4.4$ | $-26 \pm 2.2$ | 0,047 |
| S100A5-SIP | 310 | $0.92 \pm 0.24$ | 1.5 [26]* | $21 \pm 20$ | $-4.1 \pm 2.5$ | -24 |
| S100A1-NMIIA | 298 | $0.49 \pm 0.0017$ | 0.5 [18]* | $0.009 \pm 0.005$ | $36 \pm 0.34$ | -82 |
| S100B-MDM4 | 310 | $0.52 \pm 0.039$ | 1 [23] | $0.71 \pm 0.036$ | $-186 \pm 142$ | 150 |
| | | $0.53 \pm 0.038$ | | $0.63 \pm 0.081$ | $111 \pm 148$ | -148 |
| S100B-RSK1 | 298 | $0.49 \pm 0.0086$ | 0.5 [27] | $12 \pm 1.4$ | $-32 \pm 1.4$ | -60 |
| S100P-p53 | 310 | $0.28 \pm 0.0053$ | Previously unknown | $1.7 \pm 0.19$ | $-63 \pm 1.9$ | 29 |

\*These interactions were measured with a different S100 paralog

### Specificity map of the S100ome

The 20 S100 paralogs, whose interactions were studied here, represent almost the complete human S100ome [13]. It is a Chordata-specific, evolutionary young protein family, and despite the fact that they exhibit moderate sequence similarity, they are structurally very similar owing to their small size (~100 residues) and conserved fold (including two consecutive EF hand motifs) (Fig S16.). Due to this reason, their phylogenetic analysis generally does not lead to unambiguous results [29,30]. Applying different parameters during the analyses resulted in varied grouping of the human S100ome, moreover only a few clades received statistical supports (see our analyses in Fig S17.). Because of these ambiguities of the phylogenetic analyses, a phenotypic screening and analysis could provide a more reliable grouping and could reveal functional similarities among the paralogs of the protein family of interest beside the sequence-based genealogies. For such purpose, we decided to create a robust phenogram [31], representing the functional relationships within the human S100ome, using hierarchical clustering (UPGMA) [32]. This analysis separated the S100ome into two groups, in which the first group contains S100 proteins generally lacking significant interactions (termed here as “orphan” S100 proteins) and the second group comprises generally good binders (termed here as “promiscuous” S100 proteins) (Fig 4.). While promiscuous S100 proteins showed significant binding to at least a few (4-5) of the tested interaction partners, orphan S100 proteins showed either no sign of partner binding, or a weak binding to a single partner.

**Fig 4.**
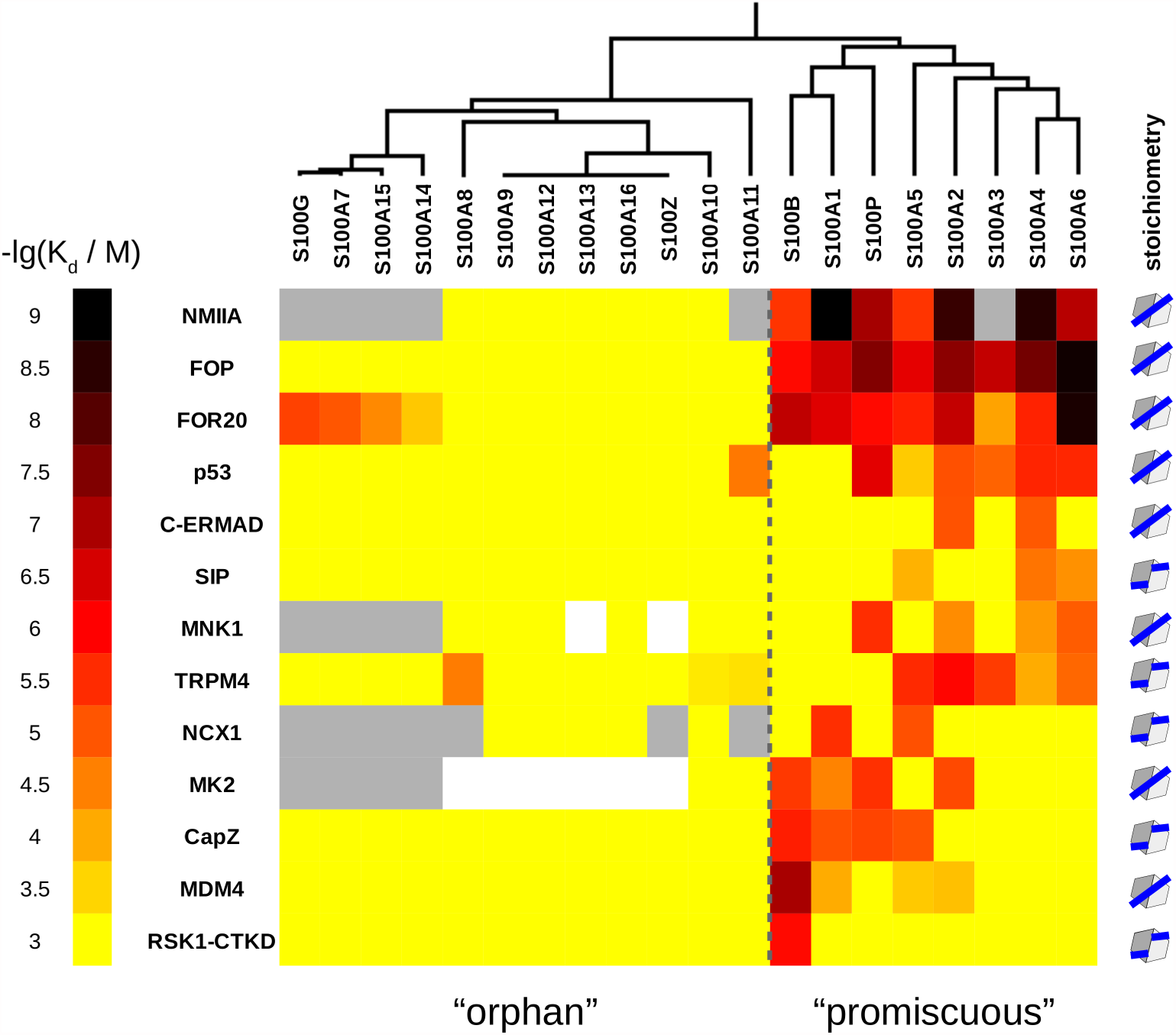
The determined dissociation constants are depicted as a heatmap representing the specificity-map of the S100ome. Hierarchical clustering, based on functional relationships, divided the S100ome into two different groups, one of them consists of low(er) specificity and/or more promiscuous S100 proteins (‘promiscuous’), while the other one contains high(er) specificity and/or less promiscuous members of the family (‘orphan’). White and grey fields indicate non-determined interactions and cases with experimental artifacts, respectively. Stoichiometry (1:1 or 2:1 to 1 S100 dimer) is also shown for all ligands at the end of each rows.

## Discussion

### Competitive FP as a potent tool to measure high-throughput macromolecular interactions

Although numerous HTP, semi-quantitative approaches are available and many low-throughput but highly accurate methods exist to measure PPIs, reliable, quantitative HTP methods are scarce in the literature. On the one hand, direct FP assay can be performed in large scale in multi-well plates which makes it an ideal method for rapid interaction screening, however, it has the serious limitation of chemical labeling that can perturb the binding measurement. Competitive FP, on the other hand, shares the same properties but without any possible interference from the labeling dye. Moreover, it provides comparative results to other, orthogonal, usually low-throughput, label-free biochemical assays, such as ITC or SPR measurements [33]. In summary, competitive FP assay is robust and HTP, thus it is a valuable tool for screening macromolecular interactions involving linear peptide motifs, RNA/DNA oligonucleotides or fluorescent small molecules [34,35].

### Evidence for functional redundancy within the S100 family and possible functions of the orphan group

S100 proteins are usually considered as ‘sticky’, relatively low specificity proteins [15], however no systematic study has been performed to make a specificity map involving the complete S100ome. Usually, the tested S100 proteins only covered the closest relatives (e.g. S100A2, S100A4, S100A6, S100B, S100P), and the results often showed redundant bindings [19,27,36–38]. Based on functional clustering, we have revealed here that the S100ome can be separated into two groups. The minor group of 8 members includes promiscuous paralogs, with a clear sign of functional redundancy. However, this does not mean that they do not have specific interactions (for example, RSK1 is highly specific partner of S100B) In contrast, the major group consists of 12 members without a clear binding preference. The function of this orphan group on the molecular level is less defined, although all S100 proteins (with the exception of S100A10) are at least involved in calcium homeostasis. They may represent intra- and perhaps also extracellular calcium ion buffers (without specific interaction partners) or they can have highly specific, yet undiscovered interaction partners [39]. As an example, S100A10, the only S100 protein without a functional EF hand motif, can mediate a very high affinity, and rather specific interaction with annexin A2 [37]. As a third scenario, it is still possible that there is functional redundancy within the orphan group, but our knowledge about S100 interaction partners is more limited in this group compared to the promiscuous group. Moreover, the present study covered only S100 homodimers (and the S100G monomer), although some S100 proteins can form heterodimers [40]. As an example, the S100A8/A9 (both coming from the orphan group) can form a functional heterodimer with known interaction partners [41].

### Function-based examination of relationships within the S100ome complements phylogenetic analysis

The phylogenetic analyses of the human S100ome resulted in rather ambiguous genealogies, likely due to the young age of the protein family (Fig S17.). Nevertheless, the clade including S100A2, S100A3, S100A4, S100A5, S100A6 was supported with high statistical values in all analyses (Fig S17.), similarly as it had also been found by others [29,30]. Our functional analysis has revealed that all members of this clade belong to the same subset of the promiscuous group, with a greatly similar functional profile. However, the phylogeny of the rest of the S100ome is supported with lower statistical values. Therefore, we suggest that in such scenarios, function-based phenotypic clustering can complement the information obtained from pure sequence-based phylogenetic analysis [42]. In our case, the S100 family can be divided, relatively unambiguously, into two bigger clusters, thus giving a more robust classification. Mapping the specificity and clustering of the S100ome contribute to the better understanding of this vertebrate-specific Ca^2+^-binding protein family. An implication of the functional redundancy defined hereby is a possibility that a function-based combinatorial S100-biomarker strategy may be more effective than detecting individual proteins.

## Materials and methods

### Expression and purification of S100 proteins

Protein preparations were done as described previously [43]. Briefly, the cDNAs of S100 proteins were cloned into a modified pET15b expression vector. All protein constructs were expressed in *Escherichia coli* BL21 (DE3) cells (*Novagen*) with a *Tobacco Etch Virus* (TEV) cleavable N-terminal His_6_-tag, and purified by Ni^2+^ affinity chromatography. The His_6_-tag was cleaved by TEV protease, which was followed by either hydrophobic interaction chromatography, ion exchange chromatography or size-exclusion chromatography with applying standard conditions [43]. The quality of the recombinant proteins were checked by SDS-PAGE analysis.

### Expression and purification of kinases

The kinase domains, MK2 (1-400) and MNK1 (1-465) were cloned into a variant pGEX expression vector. The kinase domains were expressed in *Escherichia coli* ROSETTA (DE3) cells (*Novagen*) with TEV cleavable N-terminal GST and a non-cleavable C-terminal His_6_-tag. The recombinant proteins were purified using Ni^2+^ and GST affinity purification. The quality of the kinase domains was checked by SDS-PAGE analysis. FP measurements were performed without cleavage of the GST tag.

### Expression and purification of recombinant peptides

The peptides FOR20 (1-48), FOP (1-48), p53 (1-60; 17-53), NMIIA (1894-1937), C-ERMAD (516-560 and 516-586) and RSK1 (696-735 and 689-735) were expressed in *Escherichia coli* BL21 (DE3) cells (*Novagen*) with TEV-cleavable N-terminal GST-tag, and purified by GST affinity chromatography. The tag was cleaved by TEV protease. After cleavage, the TEV protease and GST tag were eliminated by heat denaturation and centrifugation. The supernatant was purified by RP-HPLC using a Jupiter 300 Å C_5_ column (*Phenomenex*). The quality of the expressed peptides was checked by mass spectrometry (MS).

### Peptide synthesis

The CapZ (265-276), NCX1 (254-265), SIP (188-202), TRPM4 (129-147) and MDM4 (25-43) peptides were chemically synthesized using solid phase peptide synthesis (PS3 peptide synthesizer, *Protein Technologies*) with Fmoc/*t*Bu strategy in the case of (5(6)-carboxyfluorescein) labeled and unlabeled version. Peptides were purified by RP-HPLC using a Jupiter 300 Å C_18_ column (*Phenomenex*). The quality of the peptides was monitored by HPLC-MS.

### Determination of concentrations

Concentrations of peptides and proteins were determined by UV-spectrophotometry using the absorbance of Tyr and Trp residues. In the absence of these aromatic residues, the concentrations were calculated by using the absorbance of the compound on 205 and 214 nm [44,45].

### Fluorescence labeling

Chemically synthesized peptides (CapZ, NCX1, SIP, TRPM4, MDM4) were labeled with 5(6)-carboxyfluorescein at the N-terminus at the end of the synthesis. The recombinant peptides (p53, NMIIA and RSK1) were labeled with fluorescein-isothiocyanate at an N-terminal Cys residue using the protocol described previously [43]. C-ERMAD was labeled by Alexa Fluoro 568 C_5_ maleimide [19]. The excess labeling agent was eliminated by using Hitarp desalting column (*GE Healthcare*). The labeled peptides were further purified and separated from the unlabeled peptides by RP-HPLC using a Jupiter 300 Å C_5_ column (*Phenomenex*). The concentration of fluorescent peptides and the efficiency of labeling were determined by measuring the absorbance of the fluorescent dye and the peptides.

### FP measurements

Fluorescence polarization was measured with a Synergy H4 plate reader (*BioTek Instruments*) by using 485 ± 20 nm and 528 ± 20 nm, and 530 ± 25 nm and 590 ± 35 nm band-pass filters (for excitation and emission, respectively) in cases of fluorescein-based (former) and Alexa Fluoro 568-based (latter) measurements. In direct FP measurements, a dilution series of the S100 protein was prepared in 96 well plates (Tomtec plastics, PP0602; or 4titude, 96 well skirted pcr plate, 4ti-0740) in a buffer that contained 150 mM NaCl, 20 mM HEPES pH 7.5, 1 mM CaCl_2_, 0.5 mM TCEP, 0,01% Tween20 and 50 nM fluorescent-labeled peptide (probe). The volume of the dilution series was 50 µl, which was later divided into three technical replicates of 15 µl during transferring to 384 well micro-plates (Greiner low binding microplate, 384 well, E18063G5). In total, the polarization of the probe was measured at 8 different S100 concentrations (whereas one contains no S100 protein and corresponds to the free peptide). In competitive FP measurements, the same buffer was supplemented with S100 proteins to achieve a complex formation of 60-80%, based on the titration. Then, this mixture was used for creating a dilution series of the competitor (e.g. unlabeled peptide, or purified protein) and the measurement was carried out identically as in the direct experiment. Competitive FP measurement was executed if the fitted K_d_ value originated from the direct FP titration was below 200 µM. Table 1. shows the peptides used for direct and competitive FP measurements. The typical experimental window of an S100 interaction was found to be around 100 mP (polarization). However, some direct titration caused marginally small change in the polarization signal (10-30 mP), that we decided not to analyze further.

### Fitting of FP data

The K_d_ of the direct and competitive FP experiment was obtained by fitting the measured data with quadratic and competitive equation, respectively *[7]*. For automatic fitting, we used an in-house developed, Python-based program, called ProFit, which is available as a supplement of this article (see data availability section). The program is capable to process multiple experimental data at once, evaluate direct-competitive experimental data series pairs and estimate the variance of the deduced parameters (e.g. dissociation constants) through a Monte Carlo approach. It produces ready to use figures for publications, as well as a report sheet for evaluation.

### ITC measurements

Titrations were carried out either at 310 or 298 K in a buffer containing 150 mM NaCl, 20 mM HEPES pH 7.5, 1 mM CaCl_2_, 0.5 mM TCEP, using a MicroCal PEAQ-ITC instrument. The acquired data were fitted by PEAQ-ITC analysis software using the model “One Set of Sites” for most of the experiments, however for S100B-CapZ and S100B-MDM4 this model provided unsatisfactory fits and the model “Two Sets of Sites” were applied instead. Note that we used the minimal interacting region (696-735) of RSK1 instead of the larger construct (689-735), which was used in the direct FP assay.

### Bioinformatical analysis

For the phylogenetic analysis, the human S100 protein sequences were aligned using ClustalW [29] (gap open penalty 10 and gap extension penalty 0.1 for pairwise alignment, gap open penalty 10 and gap extension penalty 0.2 for multiple sequence alignment, BLOSUM weight matrix), Kalign [46], MAFFT [47] (E-INS-i strategy, BLOSUM62 scoring matrix, 1.53 gap opening penalty without offset value), Muscle [46], Prank [46] and T-Coffe [46] algorithms. Gaps were replaced by ambiguous residues (question marks) before the beginning and after the end of each sequence in the raw sequence alignment to avoid the over-interpretation of the highly variant tail extensions in the further analysis. Phylogeny was conducted with RaxML GUI [48]. Evolutionary history was inferred using maximum likelihood algorithm with ProtGamma and LG as substitution model and substitution matrix, respectively [49], with 10 runs and 1000 bootstrap replicates. For the mapping of functional relationships and clustering, the dendrogram from the S100ome data set was constructed using the unweighted pair-group method with arithmetric average (UPGMA) method [32] based on the Euclidean distance using the PAST software [50].

## Acknowledgement

We would like to thank Dániel Knapp form the Department of Plant Anatomy, ELTE for the help in bioinformatical analyses. This work was supported by the National Research Development and Innovation Fund of Hungary (K 119359 to L.N. K120391 to J.K.). M.A.S. and G.G. were supported through the New National Excellence Program of the Hungarian Ministry of Human Capacities (ÚNKP-18-2 and ÚNKP-18-3, respectively). We also acknowledge the FIEK_16-1-2016-0005, VEKOP-2.3.3-15-2016-00011 grants and Project no. 2018-1.2.1-NKP-2018-00005 implemented with the support provided from the National Research, Development and Innovation Fund of Hungary, financed under the FIEK_16, VEKOP-2.3.3-15-2016 and 2018-1.2.1-NKP funding schemes, respectively. G.G. was supported by the Post-doctorants en France program of the Fondation ARC.

## Data availability

The computer code produced for this study is available in the following website.

- ProFit: GitHub (https://github.com/GoglG/ProFit)

## Author Contributions

M.A.S. carried out the experiments, analyzed the experimental data and wrote the paper. G.G. and L.N. supervised the research, analyzed the data and wrote the paper. P.E., J.K., G.M.K. Á.L.P. and A.R. contributed by carrying out protein and peptide expression, ITC experiments and bioinformatical analyses.

